# Scalable 3D Bioprinting of Human Islets in a Pancreatic Decellularized Extracellular Matrix-Enriched Bioink for Beta-Cell Replacement Therapy

**DOI:** 10.1101/2025.06.06.658360

**Authors:** Wonwoo Jeong, Quentin Perrier, Arunkumar Rengaraj, Lori Byers, Grisell C. Gonzalez, Emma Peveri, Jake Miller, Rita Bottino, Alexei V. Mikhailov, Christopher Fraker, Emmanuel C. Opara, Alice A. Tomei, Sang Jin Lee, Giuseppe Orlando, Amish Asthana

**Affiliations:** Wake Forest Institute for Regenerative Medicine, Wake Forest University School of Medicine, Winston Salem, NC, USA; Departement of Surgery, Section of Transplantation, Wake Forest School of Medicine, Winston Salem, NC, USA; Univ. Grenoble Alpes, INSERM U1055, Department of Pharmacy, Grenoble Alpes University Hospital, LBFA, Grenoble, France; Diabetes Research Institute, University of Miami Miller School of Medicine, Miami, FL, USA; Wake Forest University, Winston Salem, NC, USA; Imagine Islet Center, Imagine Pharma, Pittsburgh, PA, United States; Department of Pathology, Atrium Health Wake Forest School of Medicine, Winston-Salem, NC, USA

**Keywords:** Pancreatic islets engineering, Type 1 diabetes, Tissue-derived biomaterials, Human islets functionalization, Microencapsulation device

## Abstract

Allogeneic cell transplantation such as beta-cell replacement for treatment of type 1 diabetes (T1D) is constrained by poor graft survival and functionality, immune rejection, and the lack of scalable biomanufacturing processes. Here, we engineered functional human islet constructs that replicate the physiomimetic human pancreatic microenvironment by employing a clinically-scalable 3D bioprinting system. To support human islet viability and function, we developed alginate-based bioinks incorporating human pancreatic decellularized extracellular matrix (dECM). These bioink formulations were optimized for shear-thinning properties for extrusion of human islets, as well as selective permeability that supports nutrient and therapeutic molecule exchange. Extrusion-based printing parameters were refined to minimize shear stress-induced damage to human islets. The resulting bioprinted pancreatic constructs demonstrated robust structural integrity, high human islet viability (>85%), and long-term glucose-stimulated insulin secretion (GSIS) over a 21-days *in vitro* culture period, even at a high islet packing density (10,000 islet equivalent/mL) while free islet controls displayed a significant functional decline. The higher performance of bioprinted islets maybe attributable to the supportive 3D dECM-rich microenvironment mitigating culture-induced stress by recapitulating the islet pancreatic niche. This scalable 3D dECM-alginate bioprinted platform represents a new advanced functional material for advancing clinically translatable bio-artificial pancreas therapies for T1D.

## 1. Introduction

Allogeneic therapeutic cell transplantation is emerging as a promising strategy in regenerative medicine, offering scalable, off-the-shelf solutions for a wide range of clinical applications, as demonstrated by encouraging outcomes from recent clinical trials across musculoskeletal, cardiovascular, and immune-mediated disorders (1–3). Allogeneic islet transplantation offers a potential superior regenerative medicine treatment for type 1 diabetes (T1D) than exogenous insulin supplementation aided by biomedical devices (4–7). However, the widespread clinical application of islet transplantation is hampered by significant challenges, including the need for chronic immunosuppression to prevent rejection and recurrence of autoimmunity, poor islet survival post-transplantation, and limitations in delivering a therapeutic islet mass in a device with suitable size for human implantation. Effective immune-isolation is crucial to prevent allograft rejection while eliminating lifelong immunosuppression. Current strategies rely on microencapsulation in hydrogel biomaterials such as alginate, or macro-scale encapsulation devices (MEDs). While microcapsules maximize the surface area to volume ratio, thereby enhancing oxygen and nutrient transport, their dispersed nature can pose challenges for surgical implantation, retrieval (if needed), and overall biosafety of this approach (8,9). On the other hand, MEDs offer retrievability in case of adverse event, yet islets entrapped in the device can agglomerate over time, exacerbating nutrient and oxygen diffusion limitations, leading to necrosis and loss of graft function (10). Similarly, embedding islets in macroscale sheet-like configurations reduces surface-to-volume ratio, which can further limit diffusion and hinder clinical scalability. A 3D bioprinted construct for islet delivery can combine the superior diffusion characteristics of microcapsules with better practical handling and retrievability of MEDs, while having the potential for scaled-up biomanufacturing for clinical applications, which could overcome the shortcomings of current islet encapsulation approaches (11).

A porous islet-delivery device with a high surface-to-volume ratio enhances nutrient/oxygen diffusion, which can increase tissue viability in bioengineered constructs. Traditional biofabrication methods can produce porous scaffolds but have limited compatibility with hydrogel biomaterials and lack precise control over construct architecture. Their reliance on thermal, solvent, or chemical treatments can damage live cells, necessitating post-fabrication cell seeding, which often leads to non-uniform cell distribution and poor attachment. Manual fabrication and assembly further limits reproducibility, hindering commercial scalability, clinical application, and regulatory compliance. Among current technologies, 3D bioprinting uniquely enables the precise, reproducible fabrication of large tissue constructs, making it particularly suitable for β cell replacement approaches. Despite its advantages, existing islet bioprinting strategies have yet to comprehensively address the critical limitations of islet delivery for T1D patients. Most bioprinted constructs are unable to simultaneously provide islet support, immunoprotection, printability, mass transport, and construct structural stability that are critical parameters necessary for therapeutic efficacy (12). For instance, a construct bioprinted entirely (or with high proportion) of decellularized extra-cellular matrix (dECM) hydrogel (13–15) or a degradable bioink (collagen (14), gelatin-based (16)) could offer cytocompatibility and vascularization potential, yet it exhibits high biodegradability *in vivo*, thus exposing islets to host immune cells leading to graft rejection. Alginate-based hydrogels have been shown to prevent direct donor-host contact but they exhibit poor printability due to their low viscosity using routine concentrations employed in islet encapsulation. Therefore, high alginate concentrations of 3-6 % (1–3) have been used to achieve a printable bioink solution viscosity, which restricted mass transport, and resulted in glucose-unresponsive or minimally-responsive islets post-printing thus significantly decreasing their glucose-stimulated insulin secretion (GSIS) functionality and their regenerative medicine applications. Human islets (HI) are highly sensitive to environmental conditions such as hypoxia and extrusion-induced shear stress, (17,18) and to date, no functional bioprinted HI construct exists that can simultaneously sustain islet viability and functionality, ensure adequate mass transport, provide structural stability, and potentially offer immune protection.

Alginate-based bioinks can offer immunoprotection but lack the biochemical cues essential for supporting optimal islet viability and function. Islet-ECM interactions are critical for regulating islet viability and function and the lack of a pancreas-specific biochemical microenvironment has been associated with limited long-term survival of islet grafts. To address this, we had previously developed a non-detergent, DI water-based decellularization protocol to produce a decellularized human pancreatic ECM powder (dECM), retaining critical matrisome components (19). Incorporating this dECM in 1.5 % alginate (Ultra-Pure Low Viscosity Mannuronate, UP-LVM) significantly improved encapsulated HI viability and function over a 58-day *in vitro* culture, providing the first evidence of the effectiveness of dECM-alginate in sustaining long-term HI functionality (19). Building upon these advancements, this study seeks to adapt this HI-supportive dECM-alginate hydrogel into a bioink suitable for reproducible fabrication of functional HI constructs through 3D bioprinting. To enhance printability without increasing the alginate concentration, we supplemented the bioink with temporary additives such as gelatin and hyaluronic acid, which optimized its viscoelastic properties for extrusion, but were washed out post-printing (20). Additionally, bioprinting parameters including extrusion pressure, speed and printed fiber diameter were optimized to maximize the construct’s diffusion efficiency and structural integrity, while minimizing the shear stress on islets during printing, ensuring optimal health and functionality of the HI-containing bioprinted construct. These optimizations are essential for producing mechanically stable constructs with islet proximity to the fiber-pore interface, that could improve nutrient transport and GSIS, unlike conventional islet hydrogels or constructs lacking microchannels and precise islet placement.

Taken together, this study pursued three key objectives: 1) the development and optimization of a bioink capable of providing a human pancreatic environment to support islets while offering potential immunoprotection, 2) optimization of bioprinting parameters tailored for HI, ensuring their viability, functionality, and the structural integrity of the printed construct, and 3) scaling up and evaluation of construct’s long-term maintenance of viability and function. For the first time, this study presents a scalable bioprinted HI construct that effectively integrates islet heath, mass transport efficiency, and structural stability. These findings represent a significant step forward in islet transplantation, offering an innovative and clinically translatable advanced functional biomaterial approach for beta cell replacement, with great potential to address key barriers to improving therapeutic outcomes for patients with T1D.

## 2. Results

### 2.1. Characterization of alginate-based bioinks for 3D bioprinting

The mechanical, rheological, and permeability properties of alginate-based bioinks were investigated to assess their suitability for extrusion-based 3D bioprinting and their potential for supporting pancreatic islet viability (**Figure 1A**). The shear-thinning behavior of the bioinks was confirmed across all formulations—LVM 1.5%, LVM 2.0%, MVG 1.1% and MVG 1.6% (**Figure 1B**). The initial viscosity at a shear rate of 1 s⁻¹ was recorded as 12.51 Pa·s for 1.5% LVM, 25.04 Pa·s for 2.0% LVM, 24.76 Pa·s for 1.1% MVG, and 43.13 Pa·s for 1.6% MVG. All bioinks exhibited shear-thinning behavior, with viscosity decreasing to 0.34-to-1.29 Pa·s at a shear rate of 100 s⁻¹, confirming their suitability for extrusion-based printing. To assess the impact of dECM concentration, 0.1–0.5 mg/mL dECM was incorporated into the 1.5% LVM bioink. The baseline viscosity of the 1.5% LVM bioink without dECM was comparable to that with 0.5 mg/mL dECM (**Figure 1C and Figure S1**). No significant change in viscosity was observed, suggesting that dECM was homogeneously solubilized and did not alter the alginate-based bioink rheological properties.

**Figure 1.**
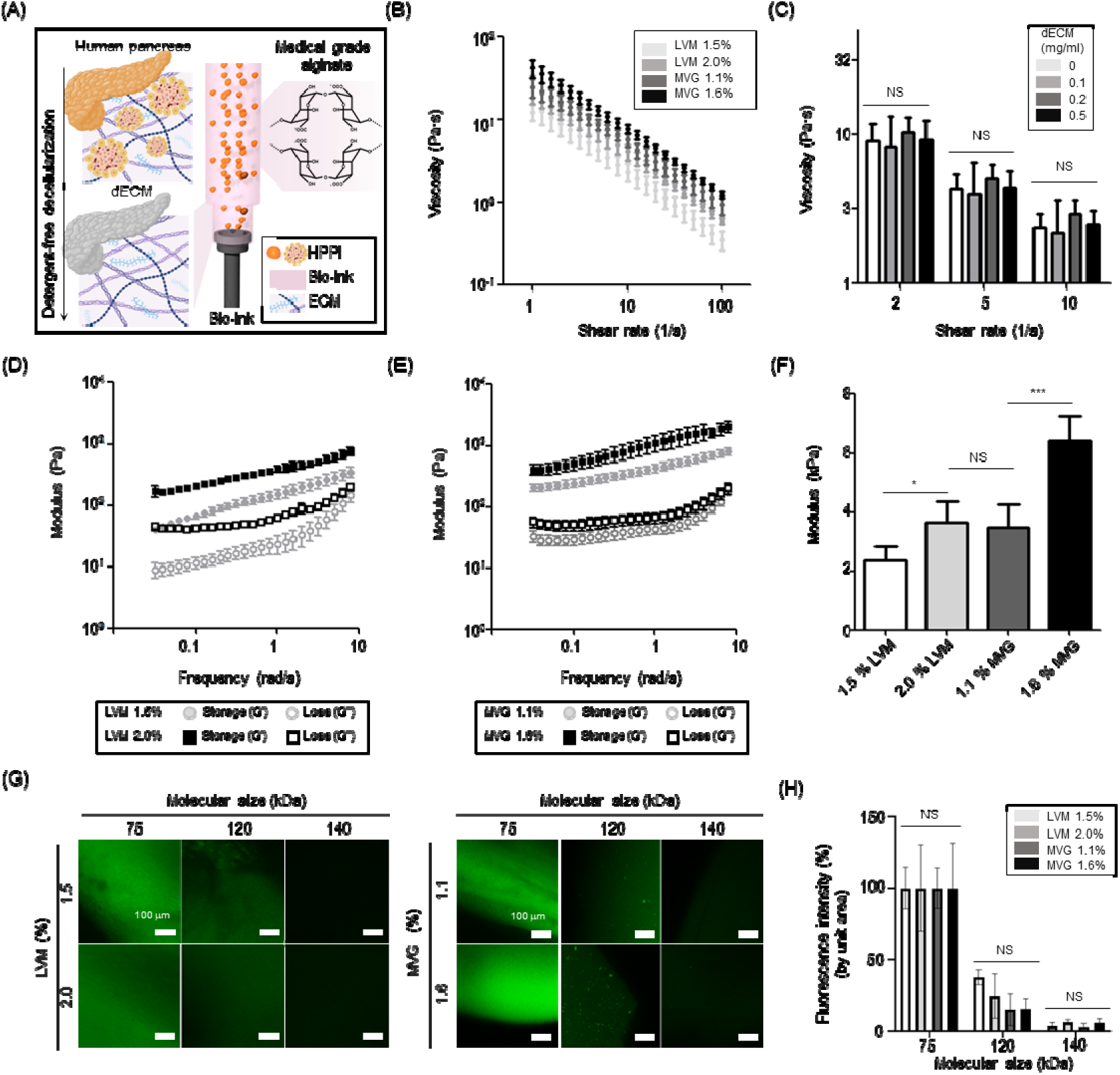
Characterization of shear behavior, compressive modulus, and molecular permeability of clinical grade alginate bioinks with pancreatic dECM. (**A**) Schematic showing preparation of pancreatic dECM-based alginate bioinks. (**B**) Shear sweep viscosity of LVM (1.5%, 2.0%) and MVG (1.1%, 1.6%) bioinks. (**C**) Viscosity of 1.5% LVM-dECM bioink containing different concentrations of dECM. (**D-E**) Frequency sweep was conducted after alginate-based bioink crosslinking with 100 mM CaCl_2_ solution. Storage and loss modulus of LVM (D) and MVG (E) were measured at 0.05 to 10 rad/s. (**F**) Compressive moduli of LVM and MVG-based bioinks. (**G-H**) Fluorescence images demonstrating the permeability of the bioinks to three fluorescent lectins with different molecular weights (MAL-1: 75 kDa, RCA: 120 kDa, and SNA: 140 kDa). Images were quantitatively analyzed for fluorescence intensity (H). Data are represented as mean with their SD. N=3, One-way ANOVA with Tukey post-hoc test *p<0.05, *** p<0.001, NS – not significant.

Viscoelastic properties of crosslinked bioinks were measured through frequency sweeps (**Figures 1D-E**). For 1.5% LVM, the storage modulus (G’) was 150.22 Pa and the loss modulus (G’’) was 23.43 Pa at a frequency of 1 rad/s (**Figures 1D**). For 1.1% MVG, G’ reached 425.06 Pa, while G’’ was 44.01 Pa at the same frequency (**Figures 1E**). Both G’ and G’’ increased with frequency across all formulations, and no crossover between G’ and G’’ was observed, demonstrating that the crosslinked bioinks maintained their elastic behavior under dynamic conditions (**Figures 1D-E**). The compressive modulus of each bioink was measured to evaluate their stiffness (**Figure 1F**). The 1.5% LVM bioink exhibited a compressive modulus of 2.36 kPa, while the 2.0% LVM reached 3.63 kPa. The 1.6% MVG bioink showed the highest modulus, which was 1.84-fold higher than that of the 1.1% MVG bioink.

Finally, permeability testing was conducted to evaluate the bioinks’ ability to allow selective diffusion of selected molecules (**Figures 1G and 1H**). After 24 h of incubation with 10 µg/mL of MAL-I (75 kDa), RCA-I (120 kDa), and SNA (140 kDa), fluorescence intensities within the alginate constructs decreased to 23.22% for 120 kDa and 4.6% for 140 kDa compared to 75kDa (100%) without significant difference among the different bioink compositions. The 75 kDa molecule (MAL-I) diffused freely through bioinks, indicating high diffusion of oxygen (0.032 kDa), glucose (0.18 kDa), and insulin (5.8 kDa). In contrast, limited diffusion of 140 kDa molecules (SNA) suggests that the bioinks can effectively restrict the movement of larger molecules, such as IgG (150 kDa) thus providing immunoisolation.

### 2.2. Optimization of islet-containing alginate-based bioprinting parameters

During the 3D bioprinting process, bioinks laden with islets or islet-like clusters are extruded through a narrow nozzle, generating shear stress that can affect the encapsulated islets. Since islets are highly sensitive to mechanical stress, it is essential to assess how printing parameters, such as extrusion pressure and printing speed, affect the morphology and viability of islets and clusters. To determine the optimal printing parameters, 1.5% LVM, 2% LVM, 1.1% MVG, and 1.6% MVG alginate-based bioinks were printed as 2D lines without cells or cell spheroids at three different extrusion pressures (30, 40, and 50 kPa) and printing speeds ranging from 10 to 120 mm/min (**Figure 2A-D**). High pressure combined with low printing speed resulted in wider fibers due to increased bioink deposition, which can hinder nutrient and oxygen diffusion from bioprinted cells. Conversely, higher speeds produced thinner fibers but increased the risk of discontinuities. Among all the bioinks tested, 1.5% LVM exhibited the narrowest printability range, likely due to its lower viscosity (**Figure 2A and Figure S2**). The optimal printing speeds for all bioinks and for each extrusion pressure were identified as the last two speeds before the printed fibers became discontinuous: 10 and 20 mm/min at 30 kPa, 30 and 40 mm/min at 40 kPa, and 40 and 80 mm/min at 50 kPa (**Figure 2A-D**). Although the bioinks displayed good printability at these parameters, further testing was required to determine their effect on cell viability and morphology.

**Figure 2.**
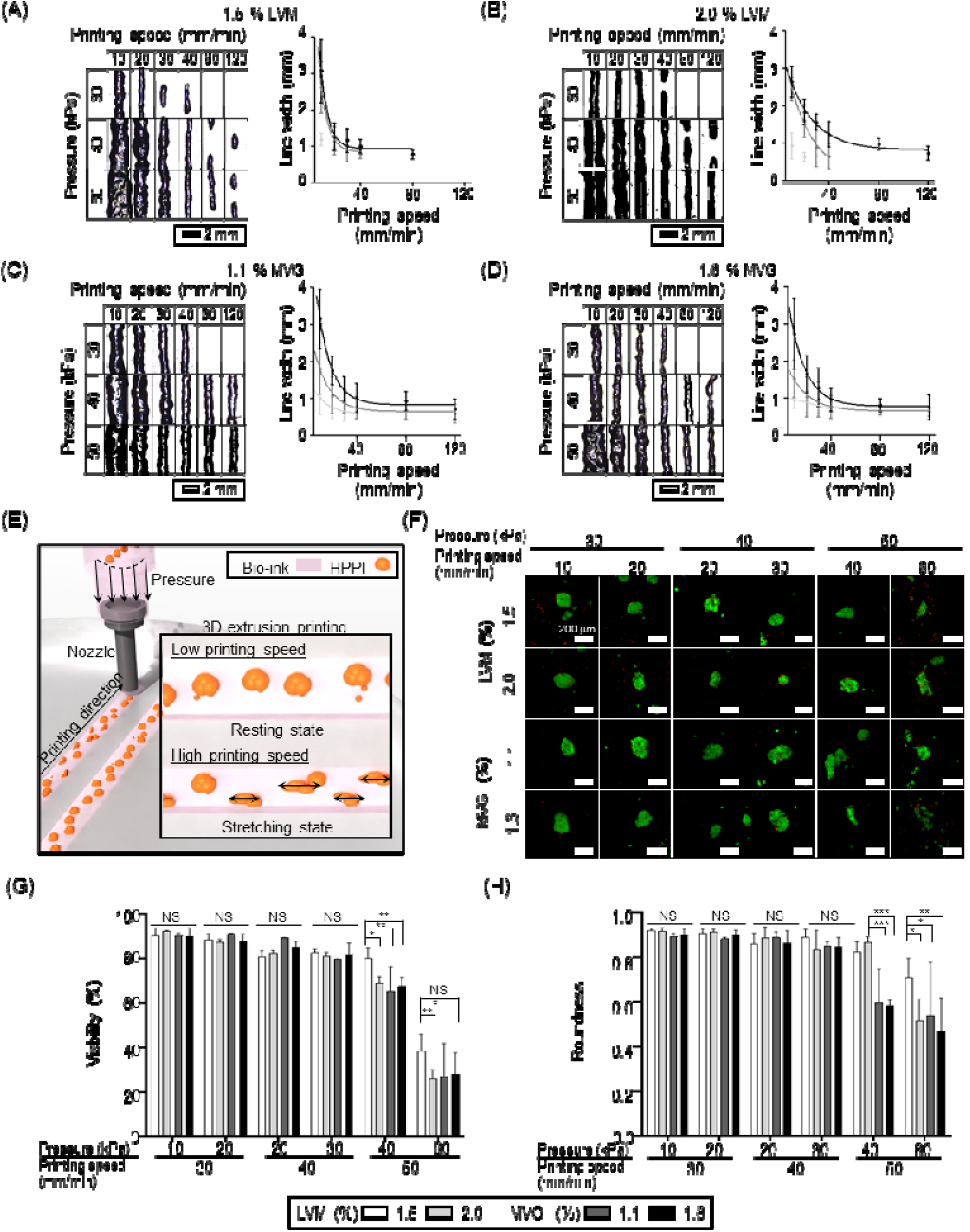
Optimization of printing parameter for alginate-based 3D extrusion bioprinting. (**A-D**) 2D line printing test results showing macroscopic images (left) and line widths (right) obtained with (A) 1.5 % LVM, (B) 2.0 % LVM, (C) 1.1 % MVG and (D) 1.6 % MVG at different printing pressures (30, 40, and 50 kPa) and printing speeds (10 – 80 mm/min). (**E**) Illustration of islet deformation due to shear stress caused by printing speed (mm/min) and extrusion pressure (kPa). (**F**) Live (green) / dead (red) confocal images demonstrating the effects of printing pressure and speed on the morphology and viability of Nit-1 clusters printed in different bioinks. Scale bar is 200 µm. Live/dead images were quantitatively analyzed for cell viability (%) (G) and cluster roundness (H). One-way ANOVA with Tukey’s multiple comparison test *p<0.05, **p<0.01, *** p<0.001, NS – not significant.

To analyze the correlation between printing parameters and clusters deformation, NIT-1 murine insulinoma cell clusters with average diameter comparable to 1 IEQ were printed using all four bioinks (**Figure 2E**). Clusters with an average diameter of 100 ± 20 µm were extruded, crosslinked, and incubated for 4 h to remove non-crosslinked additives. Fragmentation of the clusters was observed at higher extrusion pressure and printing speed, particularly for 1.6% MVG, likely due to increase in shear stress during extrusion (**Figure 2F**). Viability decreased with increasing extrusion pressure and printing speed (**Figure 2G**). At 30 kPa pressure and 20 mm/min speed, each cluster retained a viability of 88.46 ± 2.51%, maintaining structural integrity. In contrast, clusters printed at 40 and 50 kPa showed a significant drop in viability to 67.39 ± 8.05%. Only the 1.5% LVM bioink maintained a higher viability of 80.2 ± 4.32% under these conditions. Based on these results, 30 kPa and 20 mm/min were selected as the optimal printing parameters for printability while maintaining insulinoma cell cluster viability for further studies.

Similarly, extrusion pressure and printing speed influenced roundness. Clusters printed at 30 kPa and 20 mm/min maintained an average roundness of 0.90 ± 0.018. However, at 50 kPa and 80 mm/min, roundness decreased to 0.50 ± 0.16, except for the 1.5% LVM bioink, which retained a roundness of 0.71 ± 0.08 (**Figure 2H**). This indicates that the chosen parameters minimize deformation, preserving the spherical morphology critical for islet function. This conservative optimization accounted for the smaller size of NIT-1 clusters (∼100 µm), as HI are larger and more prone to shear-induced deformation. The parameters provided a baseline for future bioprinting studies involving HI. By minimizing mechanical stress during extrusion, we aimed to ensure the long-term viability and functionality of HI in printing process.

### 2.3. Optimization of 3D-bioprinted construct architecture

The mechanical and functional properties of 3D-printed constructs depend heavily on their pore size and distribution (**Figure 3**). To assess these properties, we characterized the pore fidelity and structural stability of constructs printed with different pore sizes (250 µm, 500 µm, and 1000 µm) using our alginate-based bioinks (1.5% LVM, 2% LVM, 1.1% MVG, and 1.6% MVG). Both pore fidelity (%) and shear creep/relaxation analyses were conducted to determine the optimal pore size and bioink formulation for further studies (**Figure 3A**). After crosslinking, the constructs were imaged to evaluate their pore fidelity (**Figure 3B**). Constructs printed with 1.5% LVM bioink exhibited low pore fidelity at a 250 µm pore size due to its lower viscosity. The pore fidelity for 1.5% LVM constructs was measured at 46%, 60.85%, and 82.30% for 250 µm, 500 µm, and 1000 µm pore sizes, respectively. In contrast, the 1.5% MVG construct showed well-defined pore structure and 1.07- to 1.26-fold improved pore fidelity among different pore sizes (**Figure 3C**).

**Figure 3.**
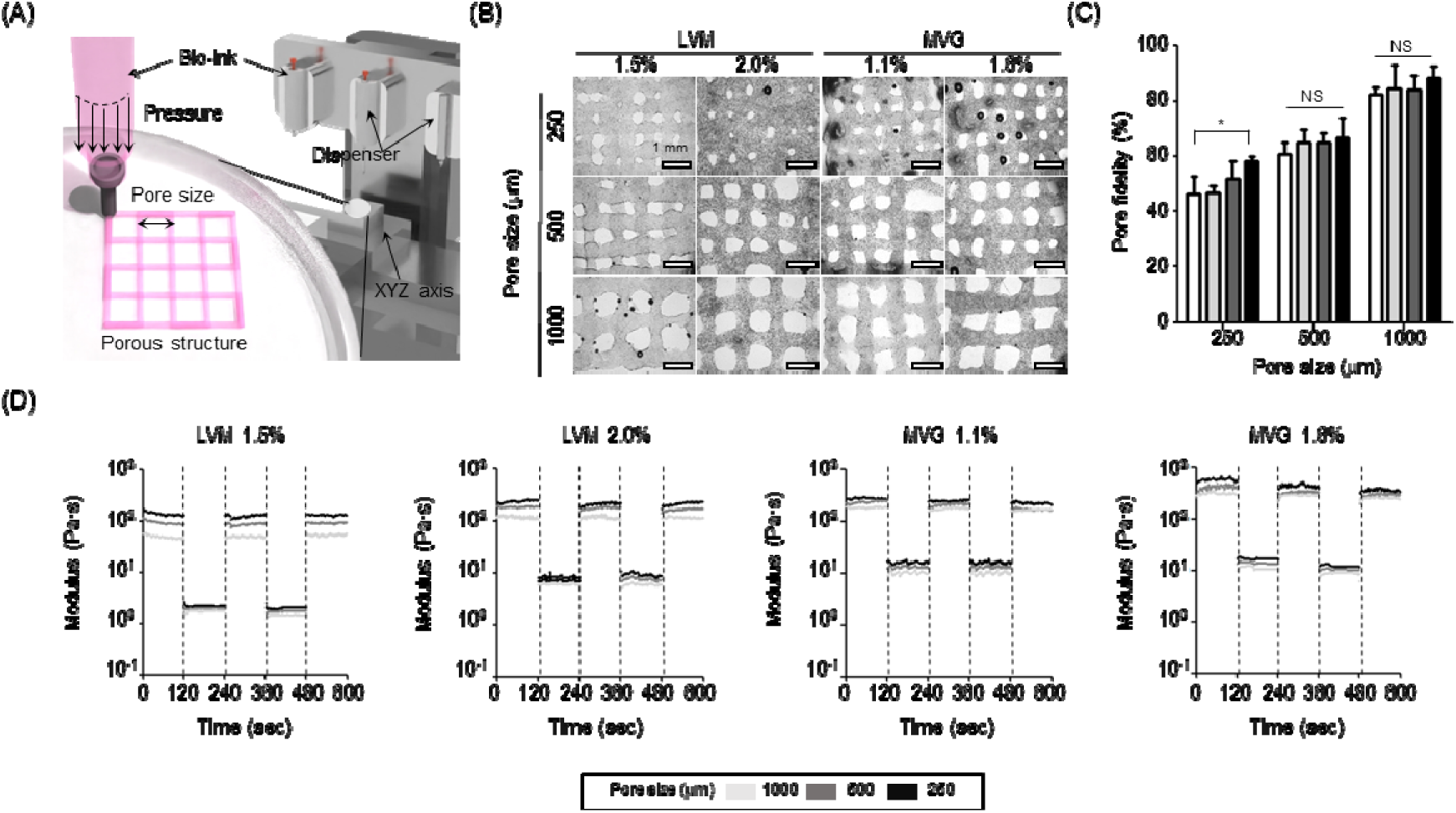
Structural and mechanical characterization of 3D printed constructs. (**A**) Illustration of the manufacturing process for bioprinting alginate-based constructs, using the Integrated Tissue and Organ Printing System (ITOP). (**B-D**) Macroscopic images (**B**) and pore fidelity assessed by quantitative image analysis (**C**) of 3D printed micro-patterned alginate constructs with different pore sizes (250 – 1000 µm). (**D**) Creep/relaxation mechanical testing of 3D printed constructs with different pore sizes, printed using the four different alginate-based bioinks. The creep/relaxation test was conducted at change of shear rates (2 /s and 50 /s). Data are represented as mean with their SD, One-way ANOVA * p<0.05, NS – not significant.

A shear creep/relaxation test was performed to simulate the structural deformation of the constructs under varying shear strain (2/s and 50/s, **Figure 3D**). Constructs were printed with a 2-layer crosshatch pattern using the different alginate-based bioinks, and the structural stability was assessed across the selected three pore sizes. The shear resistance of constructs printed with 1.6% MVG bioink and 250 µm pores was measured as 585.6 Pa·s at 2/s and 16.75 Pa·s at 50/s. This represented a 4.37-to-7.44-fold increase compared to constructs printed with 1.5% LVM bioink. These results suggest that constructs printed with 1.6% MVG bioink and smaller pores exhibit the highest shear resistance, indicating greater mechanical stability under physical stress. Overall, constructs printed with the stiffer bioink (1.6% MVG) and smallest pore size (250 µm) exhibited the highest shear resistance. Based on these observations, 500 µm pore size was chosen for future experiments, as it provided consistent printing results and good structural integrity.

### 2.4. Cytocompatibility of 3D printed constructs with NIT-1 clusters

The characterization of printability, viability, and morphology, NIT-1 clusters enabled to investigate the optimized printing parameters (20 G nozzle, extrusion pressure of 30 kPa, and printing speed of 20 mm/min) with all four bioinks (1.5% LVM, 2% LVM, 1.1% MVG, and 1.6% MVG). The alginate constructs were crosslinked with 100 mM CaCl₂ for 10 min. Live/dead staining was performed on day 0 and day 7 post-printing to assess cytocompatibility (**Figure 4A**). All printing conditions tested showed a cell viability over 80% at both time points (**Figure 4B**), indicating good cytocompatibility of the printed constructs. On day 4 post-printing, all four bioinks and free clusters control NIT-1 conditions responded to glucose stimulation, with increased insulin secretion during high glucose (HG) treatment compared to low glucose 1 (LG1), and recovery to basal insulin secretion during LG2 stimulation (**Figure 4C**). While absolute values were lower for printed NIT-1, insulin secretion normalized to the total insulin secreted during the GSIS test (sum of LG1, HG and LG2) was comparable across the different conditions (**Figure 4D**). The GSIS stimulation index (SI) was significantly higher in free clusters (SI = 2.98 ± 0.58) compared to MVG 1.1% (1.79 ± 0.26) and 1.6% (1.85 ± 0.18) but comparable to LVM 1.5% (SI = 2.66 ± 0.64) and 2.0% (**Figure 4E**), demonstrating bioink composition-dependent functional performance.

**Figure 4.**
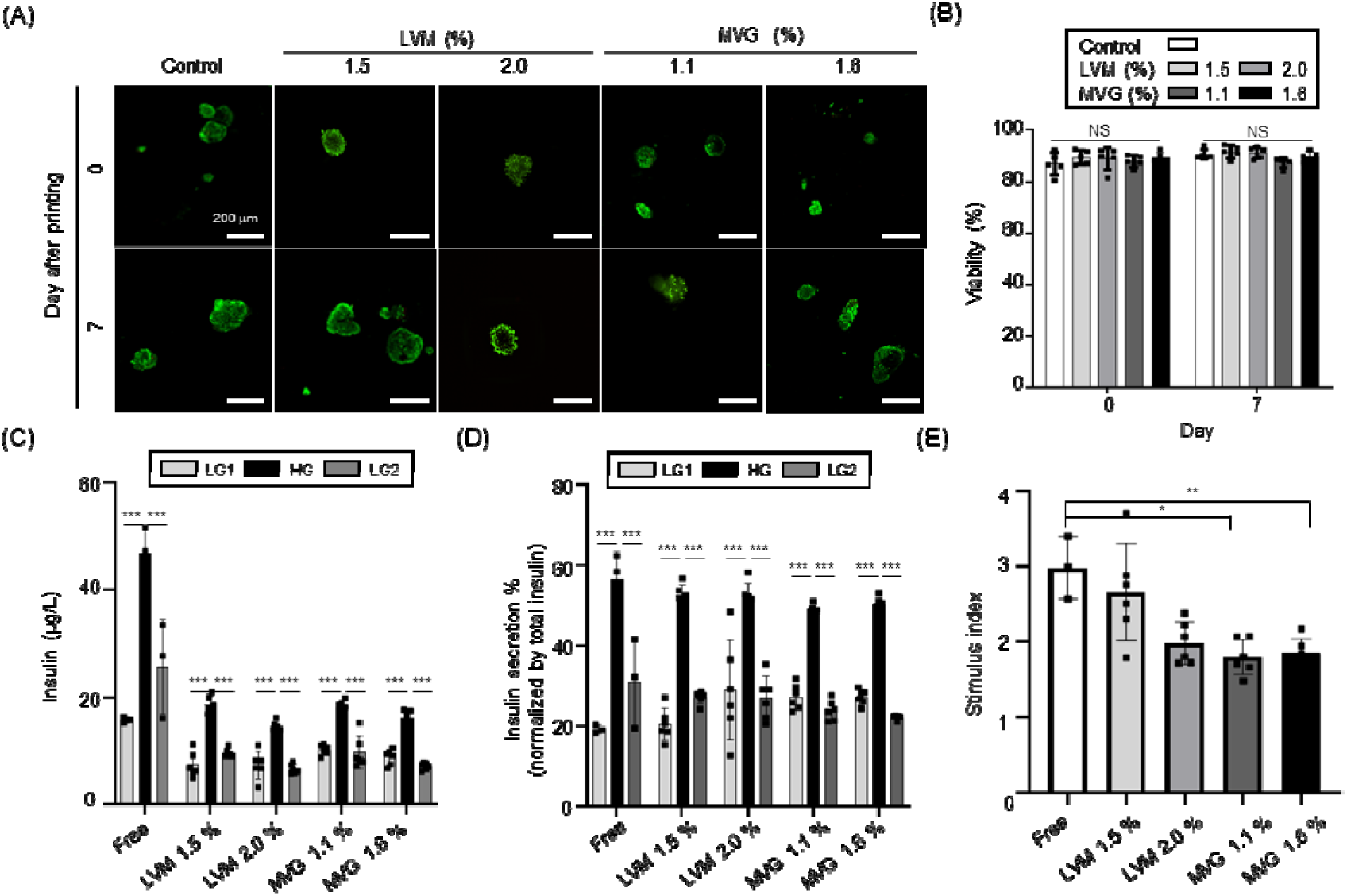
Cytocompatibility of alginate-based bioprinting with NIT-1 pseudoislets. Optimized 3D bioprinting parameters were used to test cytocompatibility of LVM (1.5%, 2.0%) and MVG (1.1%, 1.6%) based bioinks. NIT-1 murine insulionoma clusters were mixed at a concentration of 2,500 clusters/mL in the bioinks and printed at a pressure of 30 kPa and speed of 20 mm/min. (**A-B**) Representative live/dead staining confocal imaging results (**A**) and viability (%) (**B**) from 10-50 NIT1-clusters. (**C-E**) For GSIS functionality assessment, 100 free and bioprinted NIT-1 clusters were exposed to low glucose (LG1/2, 2.2 mM) for 1 hours, high glucose (HG, 16.7 mM), for 1hr, and low glucose for 1hr. Results of GSIS were presented as insulin secreted in response to glucose (**C**), insulin secretion (%) normalized to total secreted insulin during GSIS (**D**), and Stimulation index (**E**). (control; n = 3, bioinks; n = 6). Data are represented as mean with their SD, One-way ANOVA * p<0.05, *** p<0.001, NS – not significant.

Overall, this optimization study validated four alginate-based bioinks and optimized bioprinting parameters, resulting in constructs with comparable morphology and viability of non-printed NIT-1 clusters. All constructs maintained GSIS functionality, confirming the absence of mass transfer limitations. However, the functionality of MVG printed groups was lower than that of free clusters and LVM groups, likely due to the higher viscosity of MVG, which may have increased shear stress during extrusion. Further optimization of MVG bioink formulations will be required to address this issue. Given the heterogeneous size of HI and their sensitivity to shear stress, 1.5% LVM bioink was selected as the most suitable formulation for future printing of HI.

### 2.5. Bioprinting human islets with the optimized bioink and bioprinting parameters

Following the optimization of bioink formulations and printing parameters using NIT-1 insulinoma cell clusters, the next step was to apply this optimized approach for the bioprinting of HI. HI were suspended in 1.5% LVM – 0.1 mg/mL dECM bioink at a density of 2,500 IEQ/mL (HI donor information is provided in **Table S1)**. The bioprinted HI (BP HI) constructs with diameters ranging from 40 to 200 µm immediately after printing (day 0) maintained a circular morphology throughout the 7-day culture period (**Figure 5A**). This morphology was comparable to that of free HI, with average diameters of 116 ± 46 µm, suggesting that the printing process did not induce significant shear stress or morphological deformation on HI during extrusion. Additionally, the viability of both free and BP HI remained consistently above 85% at each time point analyzed and it was comparable (p>0.05, **Figure 5B**). On day 4 and 7 post-printing, GSIS functionality test was performed on free HI and bioprinted constructs. The representative insulin secretion (**Figure 5C**) and the normalized results of insulin secretion from three experiments (**Figure 5D**), highlighted the ability of HI (free and BP) to secrete insulin in response to high glucose (HG > LG1) and their ability to recover (LG2 < HG).

**Figure 5.**
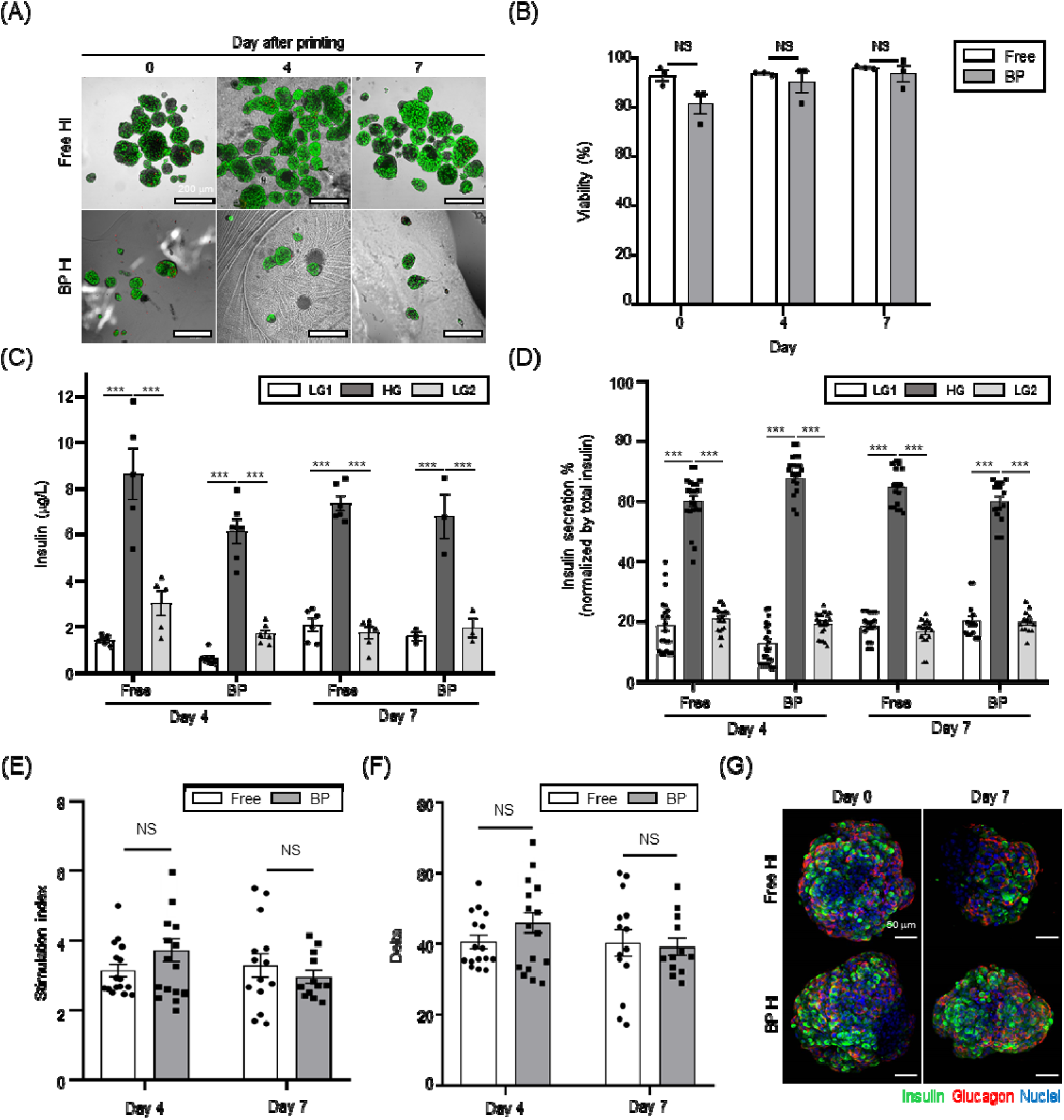
Viability and functionality of bioprinted human islets in optimized conditions. HI were printed in 1.5 % LVM - 0.1 mg/mL dECM bioink at a density of 2,500 IEQ/mL with optimized parameters (pressure 30 kPa, printing speed 20 mm/min). (**A**) Representative images of live (Syto13; green) / dead (PI; red) staining by confocal microscopy of free and bioprinted (BP) human islets (HI). 10-50 IEQ HI were used to measure the viability. (**B**) Viability of HI at day 0, 4 and 7 (N = 3). To conduct GSIS, 100 IEQ were exposed to subsequent glucose concentrations in low glucose (LG1, 2.2 mM), high glucose (HG, 16.7 mM) and LG2 (2.2mM) solutions. (**C**) Representative insulin secretion (µg/L) result from one experiment. (**D**) Insulin secretion normalized to the total insulin secreted during the GSIS at day 4 and 7. (**E**) GSIS stimulation index at day 4 and 7. (**F**) GSIS delta day 4 and 7. (**G**) Representative confocal images of insulin (green) and glucagon (red) immunostaining of free and bioprinted HI at day 0 (left) and at day 7 (right). Data are represented as mean with their SEM, N = 3 (number of HI preparations used) and n = 14-16 (number of total replicates), Two-way ANOVA with Tukey post-hoc test *** p<0.001, NS – not significant.

The GSIS SI remained comparable between free HI and BP HI at day 4 (3.15 ± 0.74 vs. 3.72 ± 1.54, p=0.444) and day 7 (3.29 ± 1.26 vs. 2.97 ± 0.65, p = 0.894, **Figure 5E**). A similar trend was observed for GSIS delta (HG insulin – LG insulin) at day 4 (40.69 ± 7.59 vs. 46.00 ± 12.69, p=0.495) and day 7 (40.30 ± 13.97 vs. 39.24 ± 8.52, p=0.995, **Figure 5F**). These results highlight that both HI groups maintained stable insulin secretion dynamics, with no significant loss in performance. Finally, the representative images of insulin and glucagon positive cells show no difference in expression of these hormones inside the HI in free HI compared to BP HI conditions (**Figure 5G**). These results aligned with the comparable viability and functionality data observed.

### 2.6. Bioprinting of high Islet density constructs

Based on the successful performance of 3D bioprinting in functional HI, a higher density of 10,000 IEQ/mL of HI was chosen for scale-up manufacturing (donor information in **Table S1**). This high-density printing was achieved without mechanical issues such as nozzle clogging. The BP HI showed no signs of islet aggregation and demonstrated a homogeneous HI distribution throughout the construct **(Figure S3)**. The morphology of the BP HI remained stable and consistent with free HI, as observed through confocal microscopy (**Figure 6A**). The viability of the BP HI (**Figure 6B**) was maintained above 80% from day 0 to day 21, with no significant changes (p>0.05), confirming the cytocompatibility of the optimized 1.5% LVM – 0.1 mg/mL dECM bioink and printing conditions. The representative insulin secretion (**Figure 6C**) and the normalized results of insulin secretion from three experiments (**Figure 6D**) showed that both free HI and BP HI were functional and responded to glucose challenges (HG > LG1 and LG2) on from day 4 to day 21. The SI of free HI decreased from 4.93 ± 0.85 on day 4 to 3.76 ± 1.07 on day 7 (p<0.05 vs. day 4) and further dropped to 2.60 ± 0.43 on day 21 (p<0.01 vs. day 7). In contrast, the SI of BP HI remained stable throughout the experiment, with values of 4.08 ± 0.93 on day 4, 3.59 ± 0.65 on day 7, and 3.68 ± 0.63 on day 21. Notably, on day 21, the SI of BP HI was significantly higher than that of free HI (p<0.01, **Figure 6E**), indicating that the incorporation of dECM components contributed to preserving long-term functionality. This result also demonstrated that the bioprinting process, even at high islet packing densities, remained stable and did not impair the functional performance of the HI over time. A similar trend was observed for the GSIS delta (**Figure 6F**). The stable SI and delta values observed in BP HI, even at high packing density, underscore the robustness of this method and highlight its potential for clinical applications in diabetes research and islet transplantation. Finally, the representative image (from one HI experiment) of insulin and glucagon positive cells show that printed HI compared to free HI after 21 days of culture exhibited more insulin positive cells (**Figure 6G**). Quantification of the proportion of beta cells after 21 days of culture showed a reduction in free HI (63.77 % vs. 47.92 % after 1 day of culture, p<0.01), whereas the proportion remained stable for the bioprinted HI (59.62 % at day 1 vs. 59.97 % at day 21, p=0.90; **Figure S4A**). Moreover, it appeared that the insulin-glucagon ratio was lower after 21 days of culture for free HI (2.13) compared to bioprinted HI (2.82, p<0.05, **Figure S4B**). The overlapping results was confirmed with a Pearson coefficient for free HI about 0.0089 (R^2^= 0.98) at day 0 and about 0.601 (R^2^= 0.95) at day 21 (p<0.001). The coefficient closed to 1 means there were overlapping between insulin and glucagon signals. Whereas the Pearson coefficient of bioprinted HI remained closed was about 0.0108 (R^2^= 0.99) at day 0 and 0.0596 (R^2^= 0.97) at day 21 with no significant difference (**Figure S4C**). These results aligned with the better GSIS results observed at day 21 for bioprinted HI compared to free HI.

**Figure 6.**
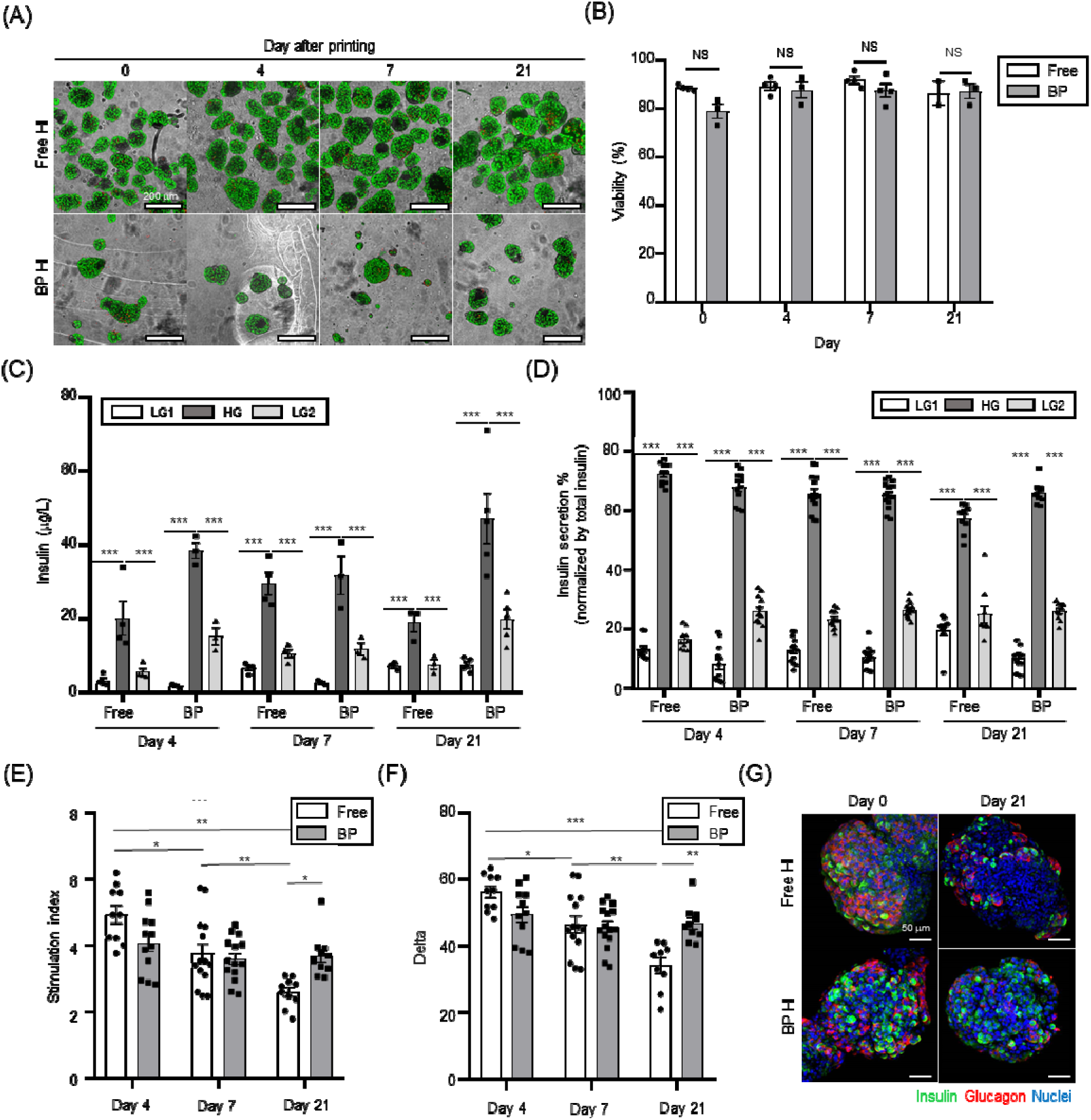
Scale-up and long-term functionality of bioprinted human islets. HI were printed in 1.5 % LVM - 0.1 mg/mL dECM bioink at a density of 10,000 IEQ/mL with optimized printing parameters (pressure 30 kPa, printing speed 20 mm/min(**A**) Representative images of live (Syto13; green) / dead (PI; red) staining by confocal microscopy of free and bioprinted (BP) human islets (HI). 10-50 IEQ HI were used to measure the viability. (**B**) Viability of HI at day 0, 4, 7 and 21 (N = 3). To conduct GSIS, 100 IEQ were exposed to subsequent glucose concentrations in low glucose (LG1, 2.2 mM), high glucose (HG, 16.7 mM) and LG2 (2.2mM) solutions. (**C**) Representative insulin secretion (µg/L) result from one experiment. (**D**)Results of insulin secretion normalized to the total insulin secreted during the GSIS at day 4, 7 and 21. (**E**) GSIS stimulation index at day 4, 7 and 21. (**F**) GSIS delta at day 4, 7 and 21. (**G**) Representative confocal images of insulin (green) and glucagon (red) immunostaining of free and bioprinted HI at day 0 (left) and at day 21 (right). Data are represented as mean with their SEM, N = 3 (number of HI preparations used) and n = 14-16 (number of total replicates), Two-way ANOVA with Tukey post-hoc test * p<0.05, ** p<0.01, *** p<0.001, NS – not significant.

## 4. Discussion

3D bioprinting has emerged as a transformative technology with the potential to address the current limitations of therapeutic cell replacement through allogeneic cell transplantation such as islet transplantation for T1D treatment by creating a structured, physiomimetic, and potentially immunoprotective environment for transplanted islets. This study reports the systematic development and validation of a scalable bioprinted HI construct, focusing on three core objectives: (1) optimization of a biomimetic bioink, (2) tailoring of bioprinting parameters specifically for HI, and (3) the evaluation of long-term HI bioprinted construct viability and function at a clinically relevant scale. Our findings demonstrate, for the first time, a bioprinted HI construct that successfully integrates sustained islet viability, efficient mass transport, structural stability, and scalability, representing a significant advancement towards a clinically translatable beta-cell replacement therapy.

The foundation of a successful bioprinting strategy lies in the bioink formulation. Using our additive-based approach, we successfully adapted both UP-LVM (1.5%, 2.0%) and UP-MVG (1.1, 1.6%) as bioinks. These hydrogels were specifically selected for their different diffusion characteristics and mechanical strengths and have been proven effective for islet immunoisolation through encapsulation, successfully preventing direct donor-host contact (21–27). However, these medical-grade ultra-pure alginates/concentrations had never been previously reported for bioprinting (28–30). All the formulations we tested exhibited essential shear-thinning behavior (**Figure 1B**), a prerequisite for extrusion-based bioprinting, which minimized shear stress during islet passage through the printing nozzle while ensuring fiber shape fidelity post-deposition. Notably, the incorporation of pancreatic dECM up to 0.5 mg/mL did not significantly alter the rheological profile of the bioink (**Figure 1C, S1**). Furthermore, permeability analysis (**Figures 1G-H**) demonstrated diffusion of molecules relevant for islet survival and function (oxygen, glucose, insulin; <75 kDa, MAL-1) while restricting the passage of larger molecules like IgG (∼150 kDa), simulated by SNA (140 kDa). This selective permeability is essential in ensuring nutrient/waste exchange and functional GSIS, while offering potential defense against humoral (IgG exclusion) immune response. This result highlighted that the powder form of the dECM, unlike hydrogels, allowed its incorporation without compromising bioink printability or altering its physical properties, seamlessly creating a biomimetic niche within any base hydrogel or bioink of choice.

HI are extremely sensitive to mechanical stress and the extrusion process would inherently subject islets to shear forces that could affect their viability and function. Therefore, optimization of printing parameters such as extrusion pressure and printing speed was essential. NIT-1 beta-cell clusters with size comparable to 1 IEQ were used as a pseudoislet model system to systematically evaluate the impact of these parameters on printed fiber width, cluster morphology and integrity, and cell viability (**Figure 2**). As expected, higher extrusion pressures and printing speeds led to increased cluster fragmentation and significantly reduced NIT-1 cell viability (**Figures 2F-G**), particularly evident with the higher viscosity MVG bioinks. This finding underscores the direct correlation between shear stress, influenced by bioink viscosity, and printing pressure and speed on cellular damage exerted during extrusion. The 1.5% LVM bioink demonstrated a wider window of cellular viability preservation, likely due to its lower viscosity imposing less shear stress. The optimized parameters (30 kPa pressure, 20 mm/min speed) achieved a high cellular viability (∼88%) while preserving cluster roundness (∼0.90), a morphological indicator often linked to islet health and function (**Figures 2G-H)**. While NIT-1 clusters are a useful surrogate, HI exhibit greater size heterogeneity and potentially higher sensitivity to printing stress. Thus, selecting conservative parameters based on NIT-1 data provided a safer starting point for HI bioprinting, prioritizing viability and structural preservation over maximizing printing speed.

Using the optimized 3D bioprinting parameters, the internal construct architecture could be precisely controlled to enhance its overall structural stability. We evaluated pore fidelity and shear resistance across different intended pore sizes (250, 500, 1000 µm) (**Figure 3**). The lower viscosity 1.5% LVM struggled to maintain fidelity at the smallest pore size (250 µm), highlighting the trade-off between lower shear stress during printing and post-printing structural definition (**Figure 3B-C**). Conversely, the stiffer 1.6% MVG bioink offered superior pore fidelity and significantly higher shear resistance, especially at smaller pore sizes (**Figure 3D**), indicating better mechanical robustness. However, considering the potential limitations of smaller pores on nutrient diffusion to densely packed islets and the consistent printing performance across formulations, a 500 µm pore size was selected as the optimal balance between structural integrity and presumed mass transport capabilities for subsequent HI experiments. This is also in corroboration with our previously published study showing that a microchannel size of ∼500 µm could provide sufficient diffusion of nutrients and oxygen in large constructs (bone; 8 × 8 × 1.2 mm^3^ and ear; 32 × 16 × 9 mm^3^) without necrosis *in vivo* (20).

NIT-1 clusters printed with the optimized parameters in all four bioinks maintained high viability (>80%) over 7 days (**Figure 4A, B**) and exhibited functional GSIS (**Figure 4C, D**). Notably, the LVM groups demonstrated a significantly higher stimulation index (SI) compared to the MVG groups, approaching the levels of free clusters (**Figure 4E**). This functional discrepancy, despite similar viability, suggested that the higher viscosity MVG bioinks might have induced stress that impacted beta-cell responsiveness, and would require further optimization. Therefore, 1.5% LVM was selected for HI printing. Transitioning to primary HIs, the optimized protocol (1.5% LVM – 0.1 mg/mL dECM bioink, 30 kPa extrusion pressure, 20 mm/min speed, 500 µm pore size) proved highly successful (**Figure 5**). Bioprinted HIs (BP HIs) maintained their characteristic morphology, exhibited high viability (>85%), and demonstrated GSIS function (SI and delta values) statistically comparable to free HI controls over the 7-day culture period (**Figures 5A-F**). Furthermore, immunofluorescence confirmed the preservation of insulin and glucagon expression within the islets (**Figure 5G**). These results demonstrated that the developed bioink formulation and optimized printing process were compatible with the sensitive human islets and effectively supported their health and function during 7-day-long *in vitro* culture.

A critical bottleneck for clinical translation is the ability to bioprint a large, therapeutically relevant islet dose in suitable device size for human implantation while maintaining long-term function. We successfully scaled up the process to print HIs at a high density (10,000 IEQ/mL) without nozzle clogging or islet aggregation, demonstrating process robustness (**Figure S3**). Remarkably, these high-density BP HI constructs maintained excellent viability (>80%) for 21 days in culture (**Figure 6B**). However, the most notable observation was the divergence in long-term function: while free HI showed a significant decline in GSIS SI by day 21, the BP HI maintained stable and robust GSIS functionality throughout the 21-day period, resulting in significantly higher GSIS SI compared to free HI at the final timepoint (**Figure 6E**). Similar trends were observed for GSIS delta (**Figure 6F**). This superior long-term HI functional preservation in bioprinted constructs was corroborated by immunofluorescence, which revealed a stable beta-cell proportion and insulin/glucagon ratio in BP HI, compared to significant beta-cell loss and altered hormonal ratio in free HI cultured under standard culture conditions for 21 days (**Figure 6G, S4A-B**). This enhanced preservation of HI function suggested that the 3D microenvironment provided by the 1.5% LVM-dECM bioink could confer significant advantages over conventional suspension culture. The structural support, combined with biochemical cues from the dECM components, likely mitigated culture-induced stress, which resulted in preserved beta-cell identity and function over extended periods, similar to the observations of our previously published study (19). This finding was supported by Kim et al. (14) bioprinted human islets with a porcine-derived pancreatic dECM-based bioink. Their preliminary data showed viable islets post-printing, but no functional data were provided. Further, Wang et al. (13), bioprinted (via stereolithography) rat islets (2,500 IEQ/mL) in 5% hyaluronic acid methacryloyl enriched with 10 mg/mL of porcine-derived dECM. They observed improved viability and functionality of the bioprinted islets *in vitro*, with higher insulin and glucagon content, and confirmed *in vivo* with diabetes reversal post-implantation. These studies combined with our results highlighted that dECM may play a crucial role in achieving optimal islet functionality.

The minimum therapeutic islet dosage for a patient is estimated to be about 5,000 IEQ/kg, which corresponds to approximately 350,000 IEQ for a 70 kg patient (6). Increasing the islet density from 2,500 IEQ/mL to 10,000 IEQ/mL made this approach clinically relevant by reducing the required volume of bioink from 140 mL to 35 mL. These high-density constructs can be printed into three implantable grafts measuring 4.18 cm per side, 2.0 mm thick, with each graft containing around 120,000 IEQ. This modular approach facilitates clinical scalability and offers surgical flexibility. Importantly, we demonstrated long-term preservation of viability and functionality in these dense constructs, supporting the feasibility of future islet graft “biobanking.” Such biobanks could enable centralized production, storage, and distribution workflows, streamlining clinical applications and reducing the need for immediate transplantation after islet isolation. Encouraged by these findings, the next step will involve preclinical evaluation in diabetic animal models to assess in vivo outcomes, including engraftment, vascular integration, long-term glycemic control, and host immune modulation. While the inclusion of 0.1 mg/mL pancreatic dECM enhanced functionality, further studies optimizing the dECM source, concentration, and specific ECM components may yield additional therapeutic benefits. Finally, transitioning toward GMP-compliant, scalable manufacturing processes will be critical for successful clinical translation.

## 5. Conclusion

This study successfully developed and validated a scalable bioprinting platform for human pancreatic islets. Through systematic optimization of bioink composition (1.5% LVM alginate with 0.1 mg/mL human pancreatic dECM) and printing parameters (30 kPa, 20 mm/min), we generated structurally stable constructs that supported high human islet viability and function during extended culture times. Crucially, the bioprinted constructs maintained superior long-term human islet functionality *in vitro* at high density (10,000 IEQ/mL) compared to standard free islet culture over 21 days. These results demonstrate the potential of this bioprinting strategy to overcome key limitations in islet transplantation by providing a supportive, scalable, and potentially immunoprotective microenvironment. This work represents a significant step towards a robust and clinically translatable tissue-engineered pancreas for the treatment of T1D.

## Materials and Methods

### Cell culture

NIT-1 mouse insulinoma cells (ATCC, VA, USA) were cultured in F-12K medium (ATCC) supplemented with 2 mM L-glutamine (Thermo Fisher Scientific), 10 %v/v fetal bovine serum (FBS, Thermo Fisher Scientific) and 1 %v/v penicillin-streptomycin (Thermo Fisher Scientific) in a humidified atmosphere at 37 °C with 5% CO₂. Cells were maintained in T75 flasks and passed every 2–3 days after reaching 80% confluence (approximately 6-7 days after seeding). Cells were then expanded into T175 flasks with tissue culture to obtain 30 million cells for clustering. For clustering, cells were seeded at a density of 30 million cells per 30 mL in the spinner flask (ReproCELL, Beltsville, MD) at 70 rpm with a magnetic stirrer (Dura-Mag, Chemglass Life Sciences, NJ, USA) kept in a CO₂ incubator. The medium was changed every other day from the 2nd day after seeding. After 4 days of culture in suspension, 50,000 IEQ of NIT-1 clusters were harvested (31). The size of the clusters was about 100-150 µm. HI were obtained from Imagine Pharma (Pittsburgh, PA, USA). Freshly isolated HI were shipped and rested for 24 h in Prodo Islets Medium Recovery (PIM(R), Prodolab, Aliso Viejo, CA, USA) in a humid atmosphere at 37 °C and 5 % CO_2_ (**Table 1**). Then HI bigger than 200 µm diameter were removed using filters (Pluriselect, El Cajon, CA, USA) and rested for 24 h in PIM Standard (PIM(S), Prodolab).

### Decellularization

Human pancreases procured from deceased donors and allocated for research purposes were obtained under an institutionally-approved protocol #IRB00028826 (Wake Forest University Health Sciences) and stored in sterile conditions at −20°C. Pancreases were collected from adult donors with a BMI (Body mass index) < 30, with no known history of diabetes. Detergent-free decellularization was performed as we previously reported (22). After removal of the peripancreatic tissue and all visible vascular structures, the tissues were cut into 1 cm^3^ pieces. To wash debris, tissue was stored in sterile water in a shaker at 4°C and 200 rpm for 24 h. The next day, a DNase solution treatment was applied for 6 h at 37°C and 100 rpm. The tissue was then washed by shaking at 4°C and 200 rpm for 18 h. At last, the tissue was transferred into a TRIS-EDTA solution and shaken at 4°C and 200 rpm for 24 h. The tissue was rinsed with sterile water and froze (−80°C). The decellularized tissue was then lyophilized and cryomilled. The decellularized tissue that met DNA (< 50 ng/mg) and endotoxin (<0.05 EU/mg) was dissolved with pepsin-HCl for 48 h at room temperature. Neutralization was performed by adding 0.1N NaOH and 10X PBS to reach a pH of 7.4 at 4C. The dissolved dECM solution was further centrifuged, and the supernatant was subjected to a series of lyophilization and cryomilling operations to produce the solubilized dECM powder. Three organs were combined into one batch to reduce variability.

### Bioink Preparation

Clinically applicable alginate-based bioinks were formulated using PRONOVA Ultra-Pure Low Viscosity Mannuronate (LVM; 1.5 and 2.0 w/v%, MilliporeSigma, Burlington, MA, USA) and PRONOVA Ultra-Pure Medium Viscosity Guluronate (MVG; 1.1 and 1.6 w/v%, MilliporeSigma), dissolved in Hanks’ Balanced Salt Solution (HBSS without calcium and magnesium, Gibco, Waltham, MA, USA). The formulation was supplemented with 0.3 w/v% hyaluronic acid (HA, MilliporeSigma) and 2.5 w/v% gelatin (MilliporeSigma). For sterilization, all bioinks were filtered using a 0.22 µm syringe filter prior to experimental use. To prepare the dECM-containing bioink, 0.1 mg/mL dECM was thoroughly mixed into the alginate-based bioink formulation before printing. Crosslinking was performed in two stages: first, the bioinks were thermally crosslinked at 4°C for 10 min to gelate the gelatin, followed by immersion in a 100 mM CaCl₂ solution in HBSS (with calcium and magnesium) at room temperature for 10 min to crosslink the alginate. After crosslinking, the bioinks were washed in HBSS without additives for 10 min twice.

### Characterization of bioinks

Rheological behavior of gelatin-crosslinked bioinks was assessed using a shear sweep test on a Discovery HR-2 rheometer (TA Instruments, New Castle, DE, USA) equipped with a 12 mm parallel Peltier plate geometry and 5 mm gap at 18 °C, simulating 3D extrusion conditions. Shear rates ranged from 0.2 to 10 s⁻ ¹. A creep/relaxation test was also performed on calcium-crosslinked 3D printed constructs with varying pore sizes, applying shear rates of 2 s⁻¹ and 50 s⁻¹ for 120 s at 1% strain. For compressive testing, cylindrical samples (8 mm diameter, 5 mm height) crosslinked with calcium were compressed at a rate of 1 mm/min using an Instron testing machine (Model 3342, Illinois Tool Works Inc., MA, USA). The elastic modulus was calculated from the slope of the stress–strain curve at 10% strain (32).

### Permeability test

Each bioink (100 µL) was two-step crosslinked in Transwell inserts (Corning) with a pore size of 12 µm and incubated for 24h in culture medium to remove HA and gelatin residues. Subsequently, the bioinks were incubated with 10 µg/mL of fluorophore-conjugated lectins of Maackia Amurensis (MAL-I, MW: 75 kDa, Vector Laboratories, Newark, CA, USA), Ricinus Communis Agglutinin (RCA-I, MW: 120 kDa, Vector Laboratories), or Sambucus Nigra (SNA, MW: 140 kDa, Vector Laboratories) for 24 h in in HBSS with calcium and magnesium. Following incubation, lectin diffusion within the alginate matrix was visualized using fluorescence microscopy (Olympus IX83, Olympus, Tokyo, JP). Fluorescence intensity was quantified by Image J.

### 3D Bioprinting system

An Integrated Tissue and Organ Printing (ITOP) system was used, equipped with a pneumatic digital precision dispense controller (ML-808GX, Musashi Engineering, Tokyo, Japan) (20). Bioinks were loaded into syringes fitted with a 20 G nozzle. The printing parameters, including printing speed (mm/min) and extrusion pressure (kPa), were adjusted according to experimental requirements. To optimize the parameters of the 3D bioprinting, NIT-1 cell clusters were suspended in the alginate bioinks at 2,500 cluster/mL and printed into a porous construct having 100 clusters. Printed morphology of the cluster was imaged by light microscopy and quantified by Image J for roundness:

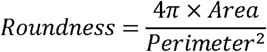

Bioprinting of HI was conducted at 2,500 and 10,000 IEQ/mL islet concentrations in the 1.5 %w/v of LVM bioink. For analysis, 100 IEQ of HI printed in 100 µL or 10 µL porous structure were used.

### Cytocompatibility test

The viability of NIT-1 clusters was assessed using a fluorescence microscope (IX83, Olympus) after staining with the LIVE/DEAD™ Viability/Cytotoxicity Kit (Invitrogen). HI viability was analyzed by confocal microscopy (Fluoview FV 3000, Waltham, MA, USA), on whole and intact islets, with 1 µM Syto13 (Thermo Fisher Scientific, Waltham, MA, USA) and 10 µg/mL of propidium iodide (Thermo Fisher Scientific). The 488 nm laser was used at 0.2% of excitation and the 561 nm lasers was used at 2.5% for excitation; the fluorescence emission was collected between 500-600 nm for Syto13 and 590-690 nm for IP. Images were acquired in z-stack with a z-step of 10 μm, a pinhole of 1 (Airy units) for all channels. Quantification was done with a macro on ImageJ software (version 8) as previously described (33). Results were expressed as percentage of viability. At least 50-100 IEQ were analyzed for each condition:

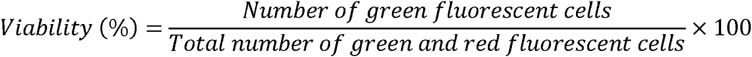

*Glucose stimulated insulin secretion* (GSIS): GSIS was performed in static incubation using Krebs-Ringer bicarbonate buffer medium (115 mM NaCl, 4.72 mM KCl, 2.56 mM CaCl_2_, 1.2 mM KH_2_PO_4_, 1.2 mM MgSO_4_, 3.89 mM NaHCO_3_, 24.4 mM HEPES, pH 7.4) supplemented with 0.2 %v/v bovine serum albumin (BSA) (w/v). Samples were initially washed in 2.8 mM glucose, followed by pre-incubation for two periods of 1 h each at 37 °C in 2.8 mM glucose. Samples were then incubated for 1 h in low glucose (2.8 mM) to measure basal insulin secretion level (LG1). Subsequently, samples were incubated for 1h in high glucose (16.7 mM) for stimulation (HG) and then incubated for an additional 1h in low glucose 2.8 mM to evaluate the return to the basal condition (LG2). All incubations were performed in a humidified atmosphere at 37 °C with 5 % CO_2_, on a rotating shaker plate. Supernatants from the different incubations were collected and frozen for later analysis. Insulin quantification assays were conducted on the supernatants from LG1, HG, LG2 conditions. The assays used mouse and human specific enzyme-linked immunosorbent assay (ELISA) kits (Mercodia, Winston-Salem, NC, USA) and absorbance was measured with the SpectraMax M5 plate reader (Molecular Devices, Sunnyvale, CA, USA). Results were expressed as the total insulin and as the percentage of secreted insulin relative to the total insulin secreted during the GSIS. The GSIS stimulation index (SI) was calculated as the ratio of insulin secreted in HG stimulation condition to the average insulin secreted in LG conditions:

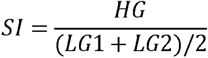

The GSIS delta was calculated with the normalized insulin secretion value:

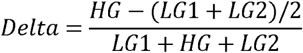

At least 100 IEQ (HI or NIT-1 clusters) were used for each replicate.

### Immunofluorescence

Free and printed HI were immunostained to assess islet cellular composition. Samples were fixed in 4 %v/v of paraformaldehyde in HBSS with calcium and magnesium overnight at 4 °C. After fixation, samples were treated with 0.2 %v/v of TritonX-100 or 10 min at room temperature. To block non-specific binding, samples were treated with 5 %w/v of bovine serum albumin (BSA, MilliporeSigma) for 1 h at room temperature. After blocking, samples were incubated with primary antibodies against insulin and glucagon overnight at 4 °C. Glucagon antibody (polyclonal rabbit anti-glucagon, PU039-UP, Biogenex) was diluted at 1:100 into ready-to-use FLEX insulin antibody solution (polyclonal guinea pig anti-insulin, IR002, Dako). After primary antibody incubation, samples were washed, followed by incubation with secondary antibodies; goat anti-rabbit Alexa Fluor 488 (A32731, Invitrogen) and goat anti-guinea pig Alexa Fluor 647 (A-21450, Invitrogen) antibodies were applied for 1 h at 1:1000 dilution. After washing, samples were stained with Hoechst 33258 (H1398, ThermoFisher). The stained samples were imaged by confocal microscopy (FV3000, Olympus).

To evaluate the co-localization of insulin and glucagon signals in immunostained samples, the Pearson’s correlation coefficient was calculated using Image J with the colocalization finder plugin. Confocal images were analyzed by selecting regions of interest (ROIs) corresponding to the islets across each z-axis focal range. The fluorescence intensity distributions of the Alexa Fluor 488 (insulin) and Alexa Fluor 647 (glucagon) channels within the selected ROIs were compared to determine the degree of spatial overlap.

### Statistical analysis

The number of total experimentations is indicated in the legend of each figure with uppercase “N” and number of total replicates with lowercase “n”. All statistical tests were performed using GraphPad (version 8.0.1). The different groups were compared by ANOVAs (or an equivalent non-parametric test if the application conditions were not met). A value is considered significant if p<0.05.

## Acknowledgement

The authors would like to thank the Center for Organ Procurement and Education (CORE), Pittsburgh, PA, which provided the donated human pancreas for isolating human islets used in this study.

## Author contribution

WJ and QP: Data Curation, Formal Analysis, Investigation, Methodology, Validation, Writing – Original Draft Preparation, Writing – Review & Editing. AK, LB, GCG: Data Curation, Methodology, Investigation. RB: Resources. EP, JM, AVM: Methodology, Investigation. CF, AAT, ECO, SJL: Resources, Supervision, Writing – Review & Editing. GO and AA: Conceptualization, Formal Analysis, Funding Acquisition, Project Administration, Resources, Supervision, Writing – Original Draft Preparation, Writing – Review & Editing

## Data availability

The authors declare that all data supporting the findings of this study are available within the paper and its supplementary information files. The associated raw data can be made available from the corresponding author upon reasonable request

## Supplementary Data

**Table S1.** Donor information of human islets.

| Supplementary Data |  |  |  |
| --- | --- | --- | --- |
| Printing condition | Age | Sex | BMI |
| 2,500 IEQ/mL | 60 | Male | 33.5 |
| 2,500 IEQ/mL | 39 | Male | 33.0 |
| 2,500 IEQ/mL | 43 | Female | 26.5 |
| 10,000 IEQ/mL | 42 | Male | 24.4 |
| 10,000 IEQ/mL | 56 | Male | 27.0 |
| 10,000 IEQ/mL | 60 | Male | 27.4 |

**Figure S1.**
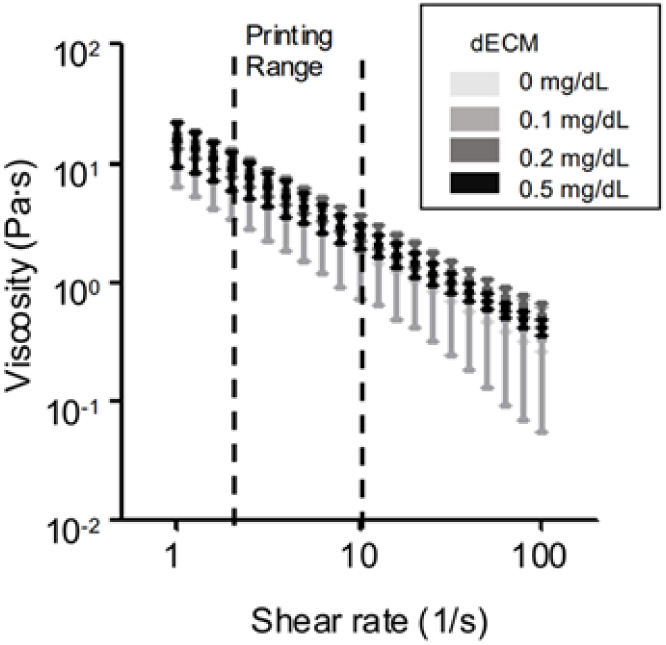
Shear sweep test of 1.5 % LVM by dECM concentration Information.

**Figure S2.**
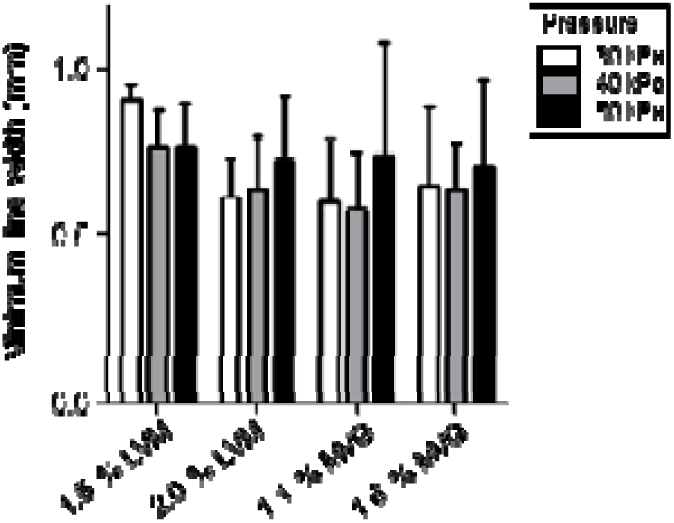
Minimum line width by pressure (30, 40, and 50 kPa) for alginate-based bio-inks.

**Figure S3.**
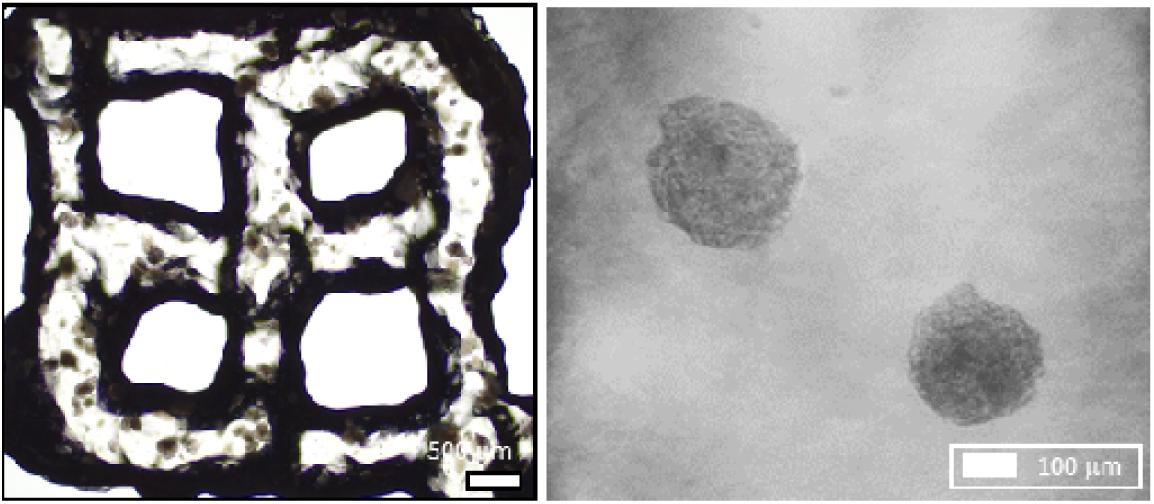
Printed construct with 10,000 IEQ/mL of HI bioink.

**Figure S4.**
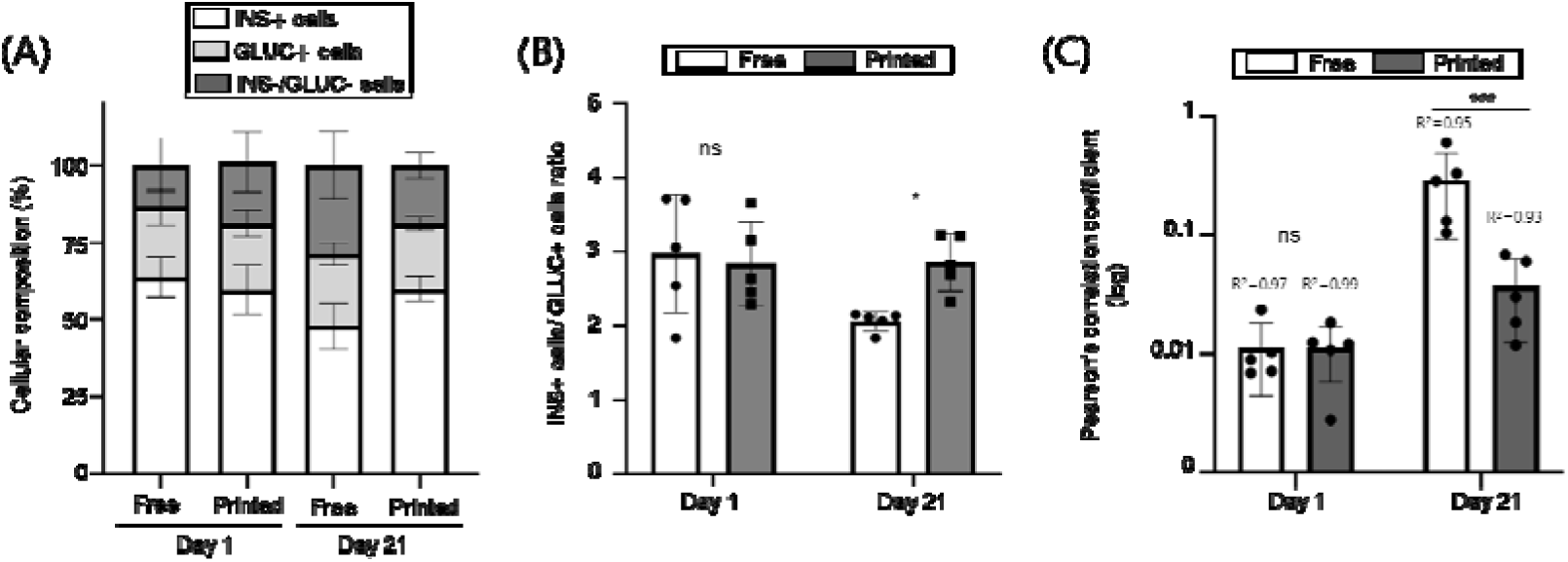
Co-localization of insulin and glucagon immunofluorescent signals in printed constructs with 10,000 IEQ/mL of HI bioink. (**A**) Cellular composition of HI as insulin positive cells (INS+), glucagon positive cells (GLUC+) and others (INS-/GLUC-). (**B**) Ratio of insulin positive cells to glucagon positive cells. (**C**) Pearson’s correlation coefficient was calculated to evaluate the co-localization of insulin and glucagon signals in immunostained samples. Data are presented as mean with their standard deviation, N = 1 (human donor) and n = 5 (fields analyzed by microscopy), Two-way ANOVA with Tukey post-hoc test *** p<0.001, NS – not significant

